# Revealing the Hidden Landscape of Public Metabolomics Data Reuse in MetaboLights

**DOI:** 10.64898/2026.05.01.722142

**Authors:** Ibrahim Karaman, Thomas Payne, Juan Antonio Vizcaíno

## Abstract

**Introduction:** Metabolomics has historically lagged behind transcriptomics and proteomics in the scale and formalisation of public data reuse. Building on earlier manually assessed studies of metabolomics data reuse, this study provides an updated characterisation of confirmed reuse in MetaboLights.

**Objectives:** This study aimed to characterise confirmed reuse of MetaboLights datasets in the published literature, quantifying its volume, reuse type, temporal trajectory, and dataset-level concentration.

**Methods:** MetaboLights dataset accessions (MTBLS#) were used to retrieve candidate publications from five literature sources via a Python-based pipeline. Candidates were manually validated and classified by primary category and scientific domain; analytical modality was assigned from dataset metadata.

**Results:** There were 274 confirmed reuse publications: Method or Tool Development (64.6%), Secondary Biological Analysis (16.8%), Data Integration or Meta-analysis (13.5%), and Educational Use or Protocol Illustration (5.1%). LC-MS was assigned to 70.1% of confirmed reuse publications; cumulative counts increased over time, and dataset-level reuse was highly asymmetric.

**Conclusion:** Within the detected publications, MetaboLights datasets were predominantly reused for computational method development with secondary biological analysis being less frequent at present, underscoring the repository’s value as a resource to obtain benchmarking datasets. Identifier-based searches may miss reuse activities when MTBLS identifiers are not consistently cited, reinforcing the community imperative for standardised dataset reporting and high-quality metadata deposition.

## 1 Introduction

Reuse of data in the public domain is now a common and defining feature of omics research. For instance, in transcriptomics, public repositories such as ArrayExpress (Athar et al., 2019) and the Gene Expression Omnibus (Edgar et al., 2002) have enabled systematic reanalysis, meta-analysis and computational benchmarking at scale using gene expression data (Rung & Brazma, 2013). In proteomics, the ProteomeXchange consortium (Deutsch et al., 2026) (including the PRIDE database) has also enabled data reuse through multiple avenues, including the development of data analysis workflows that reprocess different types of deposited MS-based proteomics data and feed results into downstream data resources (Martens & Vizcaíno, 2017; Perez-Riverol et al., 2015; Walzer et al., 2022). Across these fields, the scientific, economic and reproducibility arguments for open data reuse are well established, for instance in the context of the FAIR (Findable, Accessible, Interoperable, Reusable) data principles (OECD, 2007; Sielemann et al., 2020; Wilkinson et al., 2016). Consequently, reuse of public omics data increases the scientific return on the investments used to generate them.

Metabolomics has historically lagged behind these fields in the scale of public data sharing and formalised data reuse infrastructure. Witting observed that metabolomics data sharing is not as widespread as in transcriptomics or proteomics, attributing this partly to the technical heterogeneity of analytical platforms (LC-MS, GC-MS, NMR) and the complexity of metadata requirements (Witting, 2023). This difference in dataset availability can be quantified. By March 2026, there were 9,476 metabolomics datasets compared to 271,298 transcriptomics and 61,149 proteomics datasets among approximately 1,496,000 datasets in the Omics Discovery Index (https://www.omicsdi.org) (Perez-Riverol et al., 2017). Even when datasets are deposited, metadata gaps can render them effectively non-reusable. For instance, Harrieder et al. reported that only approximately 20% of chromatographic metadata descriptions in major metabolomics repositories were sufficiently complete to support downstream applications such as retention-time prediction (Harrieder et al., 2022).

Metabolomics data reuse also requires attention to the context of measurement. A given assay captures only a subset of the metabolome, and extraction, separation, instrument configuration, and acquisition settings influence the compounds detected and their signal response. Identification confidence and quantitative comparability further depend on supporting spectra, reference standards, calibration, processing information, and the biological and pre-analytical context of samples. These factors should be assessed before combining datasets across studies (Alseekh et al., 2021; Gouveia et al., 2024).

MetaboLights (https://www.ebi.ac.uk/metabolights) (Yurekten et al., 2024), hosted at the European Bioinformatics Institute (EMBL-EBI), is an open-access metabolomics repository endorsed by the ELIXIR European infrastructure (https://elixir-europe.org) (Harrow et al., 2021), and one of the main resources worldwide. MetaboLights is the recommended Metabolomics repository for a number of leading journals. Earlier work by Spicer and coworkers provided a manually assessed, cross-repository characterisation of metabolomics data reuse through February 2018 (Spicer, 2019). Related assessments documented limitations in metadata reporting and data-sharing practice that affect reuse (Spicer et al., 2017; Spicer & Steinbeck, 2018). Building on this precedent, the present study focuses on MetaboLights and uses an identifier-led search across five literature databases, followed by manual validation of candidate records and release of the final labelled records. A critical distinction exists between data citation—where a dataset accession appears in a reference list or data-availability statement—and confirmed active reuse, where data were downloaded and reused to produce new analytical outputs, as reported in a publication. Mixing these overstates impact and obscures the true pattern of secondary use. This report addresses that gap by providing a conservative, manually-validated characterisation of confirmed MetaboLights data reuse, covering its volume, type, temporal trajectory and dataset-level concentration.

The aim of this analysis is to characterise reuse of MetaboLights datasets in the scientific literature. Two complementary emphases guide the work: first enumerating the population of unique publications that have reused at least one MetaboLights dataset, and second identifying which MetaboLights datasets have been reused most frequently and describing the recorded characteristics of those datasets.

The scope is restricted to MetaboLights (MTBLS) in this report. Data reuse is defined conservatively: only manually-validated cases are counted. Cases where a dataset accession is mentioned without evidence that the data was actively reused and cases where the publication is the main (submission) reference for a dataset accession are excluded from this work.

## 2 Materials and Methods

Data retrieval for review and further analyses was performed in January 2026 using a Python workflow, with analysis restricted to public MetaboLights (MTBLS) datasets available at that time.

### 2.1 Search and Retrieval

In January 2026, we retrieved the list of 2,560 public MetaboLights dataset accessions (or IDs) at the time through the MetaboLights public web-service endpoint. Each accession was used to query five literature sources: PubMed Central, Europe PMC, ScienceDirect, Springer, and Wiley. The source-specific collectors used the search and retrieval facilities made available by each provider, including full-text indexes, APIs, or RSS-based searches where applicable. The search targeted explicit MTBLS accession identifiers in searchable article text; supplementary files were not systematically searched unless indexed as part of the source’s searchable full text. The search used the full list of 2,560 public MetaboLights accessions, with no additional user-defined cap on the number of records retrieved per accession from each source. Retrieval remained subject to the source-specific limits described in Table 1. Publication year followed source metadata: PMC publication date, then electronic-publication date; Europe PMC reported year; ScienceDirect cover date; and Springer/Wiley publisher dates, then RSS published and updated dates. Records manually classified as published preprints were excluded, and only candidates traceable to the January 2026 MTBLS-identifier search were retained. The example queries in Table 1 use MTBLS1 as an illustrative accession. The released collectors retrieve a single API result page or RSS feed per accession without pagination; exhaustive retrieval is therefore not guaranteed.

**Table 1.** Source-specific search interfaces, example queries, and built-in retrieval limits in the released workflow.

| Source | Interface and query | Built-in retrieval limit |
|---|---|---|
| PubMed Central | ESearch: db=pmc, term=MTBLS1 | retmax=1000 |
| Europe PMC | Search API: query="MTBLS1", resultType=core | pageSize=1000 |
| ScienceDirect | Search API: query="MTBLS1" | count=100 |
| Springer | Springer Link RSS: query=MTBLS1, sortBy=newestFirst | Entries returned in one RSS feed |
| Wiley | Search RSS: AllField=MTBLS1, content=articlesChapters | Entries returned in one RSS feed |

The released search workbook contains 4,268 candidate publication-dataset hits across its five source tabs: 1,715 from PubMed Central, 1,800 from Europe PMC, 332 from Springer, 298 from ScienceDirect, and 123 from Wiley. Candidate hits whose DOI matched a dataset’s repository-recorded primary publication DOI were removed automatically. Remaining candidate hits were merged across sources into the Combined candidate sheet used to prepare the reviewer workbook. The released collector code specifies case-insensitive DOI plus MTBLS accession for hits with a DOI, or normalized title, publication year, and accession for hits without a DOI, with source priority PMC, Europe PMC, ScienceDirect, Springer, then Wiley. The 1,239-row reviewer workbook retained 30 duplicate candidate rows retrieved from multiple sources, identified by matching DOI and MTBLS accession (29 rows), or title, publication year, and accession where no DOI was available (one row). After manual review, publication identities were deduplicated within each MTBLS accession for the final analyses, as described in the section 2.3. The released search workbook is a filtered, analysis-ready copy of the January 2026 search, covering publications dated through 2025.

### 2.2 Manual Validation

Candidate hits were manually reviewed by a single domain expert using predefined data reuse definitions. Each candidate was assessed for whether the dataset accession appeared in the context of active data use (e.g., reanalysis or integration) or just as a reference citation (mention or submission reference / original manuscript). Each confirmed reuse case was annotated by the reviewer with a primary category, sub-category and scientific domain (metabolomics, lipidomics, both, or other), based on the study context and reported analyses. Confirmed reuse was recorded in the Reviewer.Notes column using the labels Reuse, Benchmark or Reanalysis. The primary categories and sub-categories were defined based on common data reuse patterns observed during an initial screening and informed by the existing use cases in the metabolomics and broader omics data reuse literature. The primary categories are described in Table 2. The sub-category titles are presented in Supplementary Table S1. ChatGPT (OpenAI; GPT-5.1, used on 20 November 2025) was used only to assist compilation of an initial candidate list of reuse categories. It was not used to screen publications or to determine inclusion, exclusion, primary category, sub-category, or domain assignments; these decisions were made manually by the reviewer. Automated steps retrieved and merged candidate identifier-publication matches and prepared the reviewer workbook; publication-dataset pairs were deduplicated for analysis after manual screening. The domain expert manually identified original/submission reference papers that were not already recorded in MetaboLights, assessed active reuse versus citation-only mentions, and assigned the final inclusion decision, primary category, sub-category, and domain.

**Table 2.** Reviewer-assigned primary reuse categories.

| **Primary Category** | **Description** |
|---|---|
| Method or Tool Development | Datasets used to develop, evaluate, benchmark, or compare analytical methods, software tools, or computational workflows. For example, benchmarking a preprocessing or annotation workflow. |
| Secondary Biological Analysis | Datasets re-analysed to generate new biological findings beyond the scope of the original study. For example, identifying a biomarker or disease-associated feature. |
| Data Integration or Meta-analysis | Datasets aggregated, harmonised, or integrated across studies or omics layers for cross-platform, cross-study or multi-omics synthesis. For example, harmonising multiple cohorts or integrating omics layers within a single study. |
| Educational Use or Protocol Illustration | Datasets used for demonstration, training, tutorials, or workflow illustration without producing new scientific findings. For example, using an archived dataset in a tutorial or protocol. |

A publication was classified as confirmed reuse when it provided explicit evidence that one or more deposited MTBLS datasets were actively reused to produce a new analytical output. Eligible uses included reanalysis, method or tool development and benchmarking, and data integration across datasets or studies. Each confirmed reuse publication received one primary category and one sub-category reflecting its principal analytical purpose. We excluded original/submission reference papers for the dataset concerned, identifier-only mentions in references or data-availability statements, and papers without evidence that the deposited data were analysed. For example, a paper that benchmarked a method using an MTBLS data matrix was included even if it made no new biological claim; a paper that merely cited an MTBLS accession without analysing the associated data was excluded.

### 2.3 Handling Duplicates and Metadata Merging

After manual validation, records were grouped by DOI where available. A record without a DOI was linked to another record’s DOI only when they shared a PMCID associated with exactly one DOI in the reviewed corpus. Otherwise, records were identified by PMCID or, when neither identifier was available, by normalized title. Records with different DOIs, including preprint and journal versions, were kept separate; their inclusion followed the expert’s manual screening decision. For each publication–MTBLS accession pair, one record was retained. Where duplicate records had different screening decisions, a record labelled Reuse, Benchmark or Reanalysis was retained in preference to an excluded record. Confirmed publication–dataset associations were then merged with MetaboLights metadata for stratified analysis.

Publication-level summaries count each confirmed publication once, irrespective of the number of linked datasets. Dataset-level summaries count each distinct confirmed publication–MTBLS accession association.

## 3 Results

### 3.1 Data Summary

Table 3 summarises the search and manual validation results for MetaboLights. Of the 884 unique publications identified, 274 were confirmed to have reused at least one MetaboLights dataset, and 610 were excluded. Among the excluded publications, 553 were submission references for the deposited dataset (90.7%), 26 were mention-only publications without evidence of active reuse (4.3%), 14 were published preprints (2.3%), 8 were wrong citations (1.3%), 4 had no accessible full text (0.7%), 2 were labelled as duplicates (0.3%), 2 were false positives (0.3%), and 1 was a correction/erratum (0.2%). Each publication received one final screening outcome. Where records had different screening decisions, confirmed reuse took precedence; among exclusions, submission-reference or mention-only decisions took precedence over duplicate labels. Preprints and journal articles with different DOIs were treated as separate publications, and preprint exclusions followed the manual screening decisions. The full list of confirmed dataset–publication associations and the underlying summary results are provided in Supplementary Data S1.

**Table 3.** Search and validation summary across MetaboLights (MTBLS).

| Metric | Count |
|---|---|
| publication - dataset accession matches | 1,234 |
| Unique dataset accessions | 896 |
| Unique publications | 884 |
| Unique publications that are submission reference | 553 |
| Mention-only publications (no evidence of active reuse) | 26 |
| Published preprints | 14 |
| Duplicate records | 2 |
| Wrong citations | 8 |
| No accessible full text | 4 |
| False positives | 2 |
| Corrections or errata | 1 |
| Total excluded publications | 610 |
| Unique publications with confirmed data reuse | 274 |

### 3.2 Overall Reuse Patterns

Figure 1 shows the distribution of the 274 confirmed reuse publications by primary category, analytical modality, and scientific domain. Method or Tool Development accounts for 177 of the 274 unique reuse publications (64.6%), making it the largest primary category. Secondary Biological Analysis contributes 46 publications (16.8%), Data Integration or Meta-analysis 37 (13.5%), and Educational Use or Protocol Illustration 14 (5.1%). LC-MS was the most frequent publication-level modality assignment, accounting for 192 publications (70.1%), followed by NMR (37, 13.5%) and GC-MS (19, 6.9%). The remaining modality assignments were MS (other/unspecified) (15, 5.5%), LC-MS & GC-MS (8, 2.9%), LC-MS & NMR (2, 0.7%), and LC-MS & GC-MS & NMR (1, 0.4%). These publication-level modality proportions are descriptive and are not directly comparable with repository-level accession distributions; they should not be interpreted as evidence of preferential reuse. Most confirmed reuse publications were assigned to Metabolomics (252, 92.0%), followed by Lipidomics and Other (10 each, 3.6% each), and Both (2, 0.7%).

**Fig 1.**
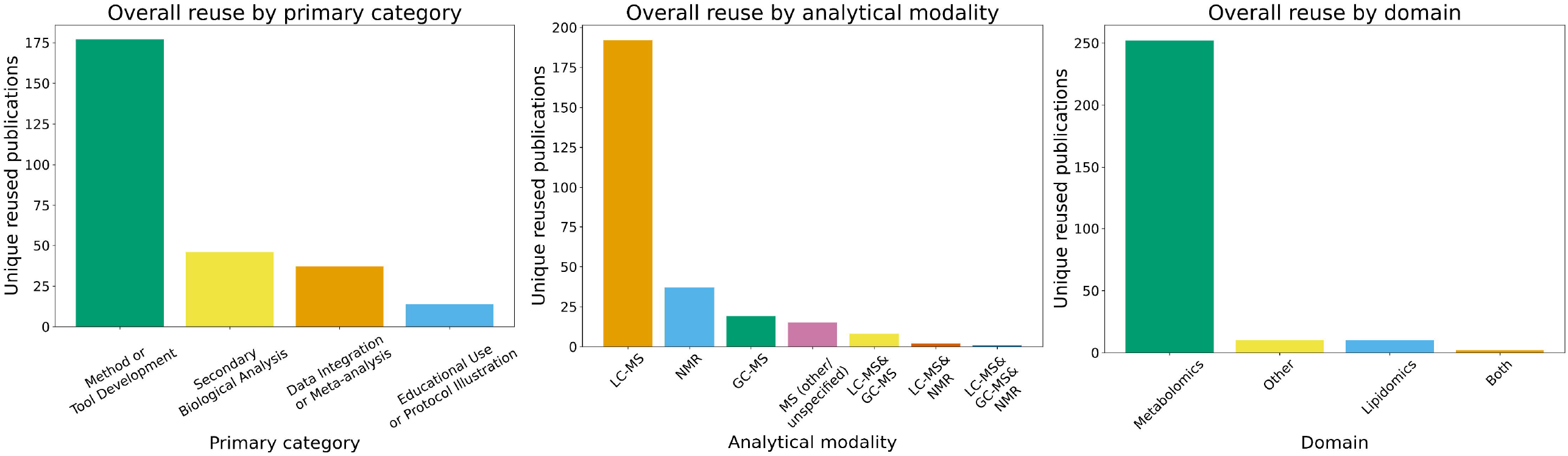
Distribution of the 274 confirmed reuse publications by primary category (left), analytical modality (centre), and scientific domain (right). Each publication is counted once in each panel. Modality was derived from MetaboLights dataset metadata, using the most frequent modality label across each publication’s linked datasets; ties were resolved deterministically by the analysis code. “MS (other/unspecified)” denotes other or unspecified MS methods, and ampersand-separated labels indicate datasets with multiple modalities. “Both” denotes metabolomics and lipidomics; “Other” denotes publications whose principal domain was neither.

### 3.3 Time-based Trends in Reuse

#### 3.3.1 Cumulative Growth

Figure 2 shows the cumulative number of confirmed reuse publications by publication year. The total exceeded 100 publications in 2021 and 200 in 2024, reaching 274 by the end of 2025. These are absolute counts of detected reuse publications; they do not establish changes in the likelihood that a dataset will be reused.

**Fig 2.**
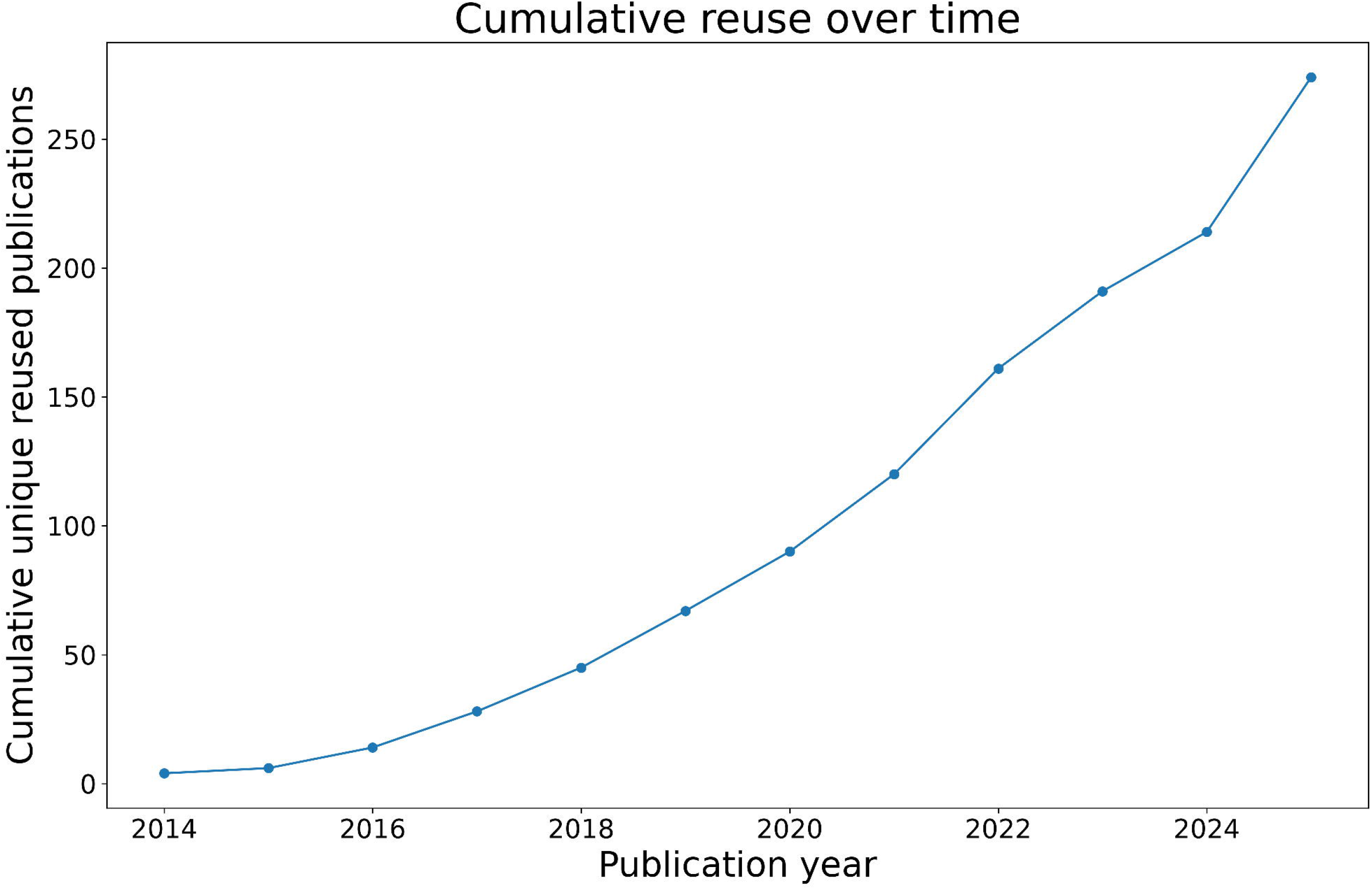
Cumulative number of confirmed reuse publications by publication year. Each publication is counted once. Counts for 2025 may be incomplete because the January 2026 search may not have captured publications that had not yet been indexed by the literature sources.

#### 3.3.2 Annual Reuse by Category

Figure 3 shows the cumulative (left) and annual (right) numbers of confirmed reuse publications by primary category. Method or Tool Development accounted for the largest annual count in every year from 2014 to 2025. The first identified publications classified as Secondary Biological Analysis and Data Integration or Meta-analysis were dated 2016 and 2017, respectively. Annual counts varied across categories, with year-to-year increases and decreases. These patterns describe absolute numbers of detected reuse publications rather than category-specific reuse rates.

**Fig 3.**
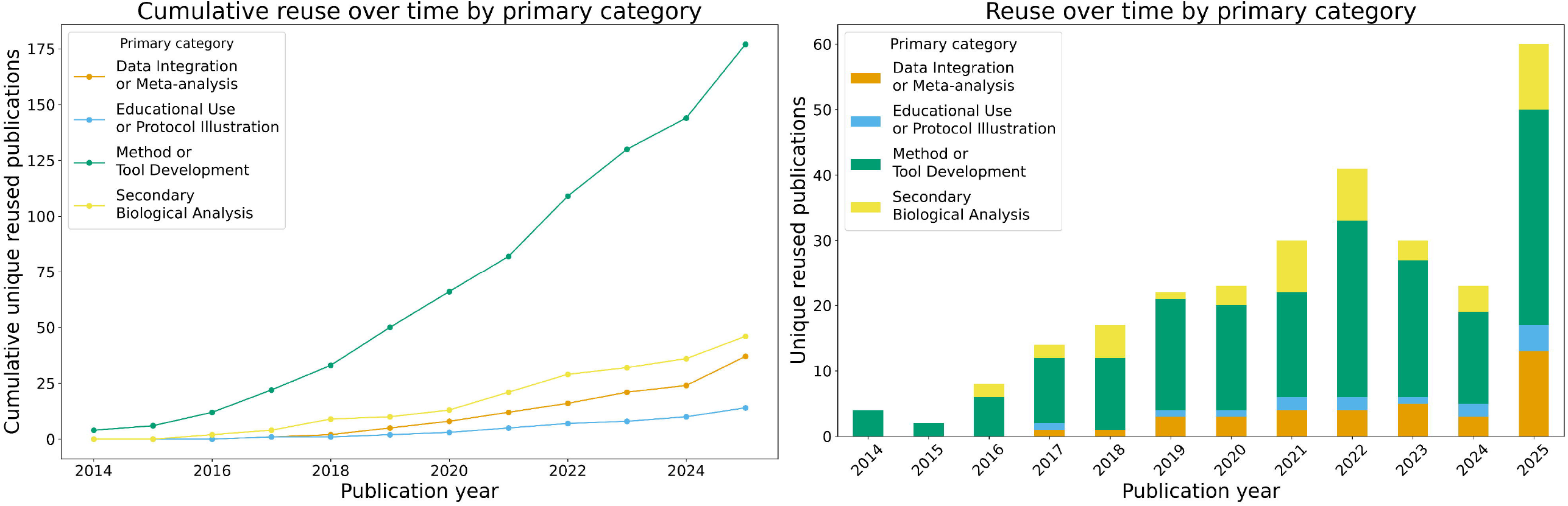
Cumulative (left) and annual (right) numbers of confirmed reuse publications by primary category. Each publication was assigned one primary category. Counts for 2025 may be incomplete because the January 2026 search may not have captured publications that had not yet been indexed by the literature sources.

#### 3.3.3 Annual Reuse by Modality and Domain

Figure 4 shows the annual numbers of confirmed reuse publications by analytical modality and scientific domain. LC-MS was the most frequent modality assignment overall, although annual counts fluctuated, reaching 45 publications in 2025. NMR contributed smaller annual counts, including six publications in 2025. Metabolomics was the most frequent domain assignment in most years, whereas Lipidomics accounted for fewer publications and appeared in five of the twelve years. These patterns describe absolute publication counts rather than modality- or domain-specific reuse rates.

**Fig 4.**
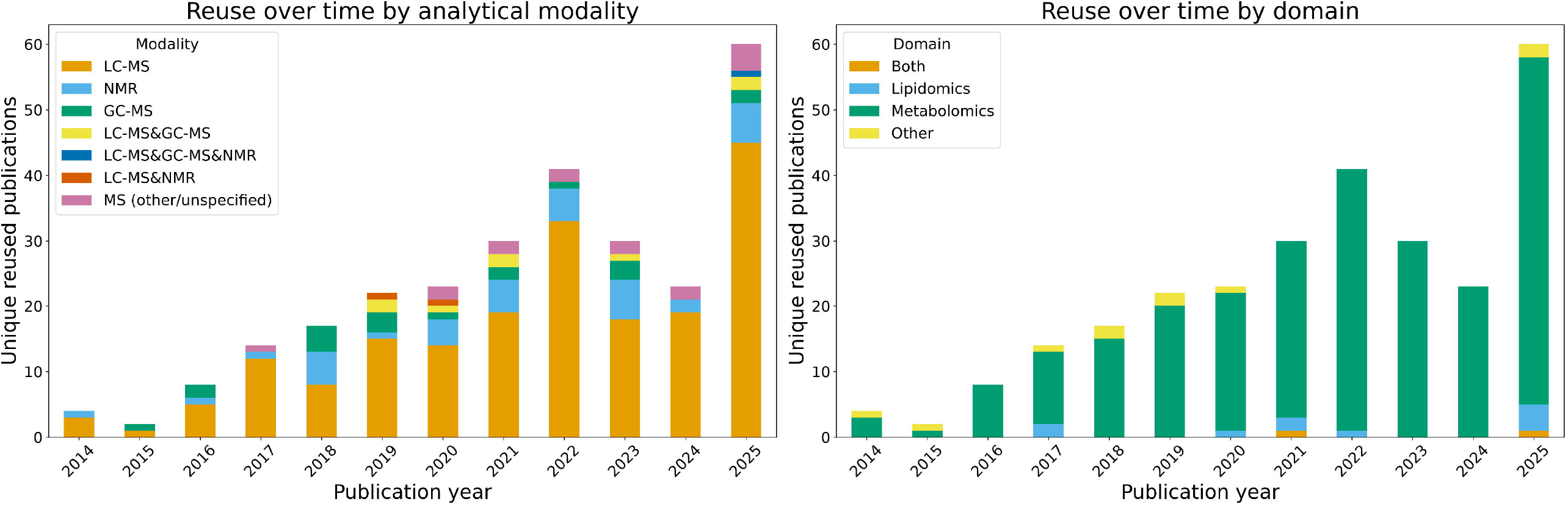
Annual numbers of confirmed reuse publications by analytical modality (left) and scientific domain (right). Each publication is counted once in each panel. Counts for 2025 may be incomplete because the January 2026 search may not have captured publications that had not yet been indexed by the literature sources.

### 3.4 Category, Modality, and Sub-category Interactions

Interactions between primary categories, analytical modalities, and sub-categories were analysed using aggregated counts of unique reuse publications. Detailed results are shown in Supplementary Figures S1-S3.

The largest category–modality combination was Method or Tool Development paired with LC-MS, accounting for 124 publications (Supplementary Figure S1). Secondary Biological Analysis shows a similar pattern but at a smaller scale. Data Integration or Meta-analysis is more focused, with most data reuse coming from multi-omics integration studies.

At the sub-category level, data reuse is concentrated in a few common tasks. Within Method or Tool Development, the main areas are Identification/Annotation, Preprocessing/Deconvolution, and Analysis Workflow. Machine Learning accounts for 15 publications. Within Secondary Biological Analysis, most data reuse relates to biomarker discovery and case-control analysis.

NMR and GC-MS are used in more specific contexts, for example mainly in preprocessing and identification tasks for NMR, and classification/regression and preprocessing tasks for GC-MS (within Method or Tool Development).

These interaction results are descriptive counts of identifier-linked reuse publications.

### 3.5 Data Reuse Focus: The Most Reused Datasets

#### 3.5.1 Highly Asymmetric Distribution

Of 273 unique MTBLS dataset accessions appearing in confirmed data reuse publications (274 reuse publications), 184 (67.4%) are associated with a single reuse publication, 73 (26.7%) from two to four reuse publications, and 16 (5.9%) with five or more. This indicates a highly asymmetric distribution of dataset-level reuse within the confirmed set.

#### 3.5.2 Top 20 Most Reused Datasets

Figure 5 shows the top 20 most reused MetaboLights datasets by total publication-dataset matches, coloured by primary category. MTBLS1, MTBLS28 and MTBLS90 lead with 10 confirmed publication-dataset matches each.

**Fig 5.**
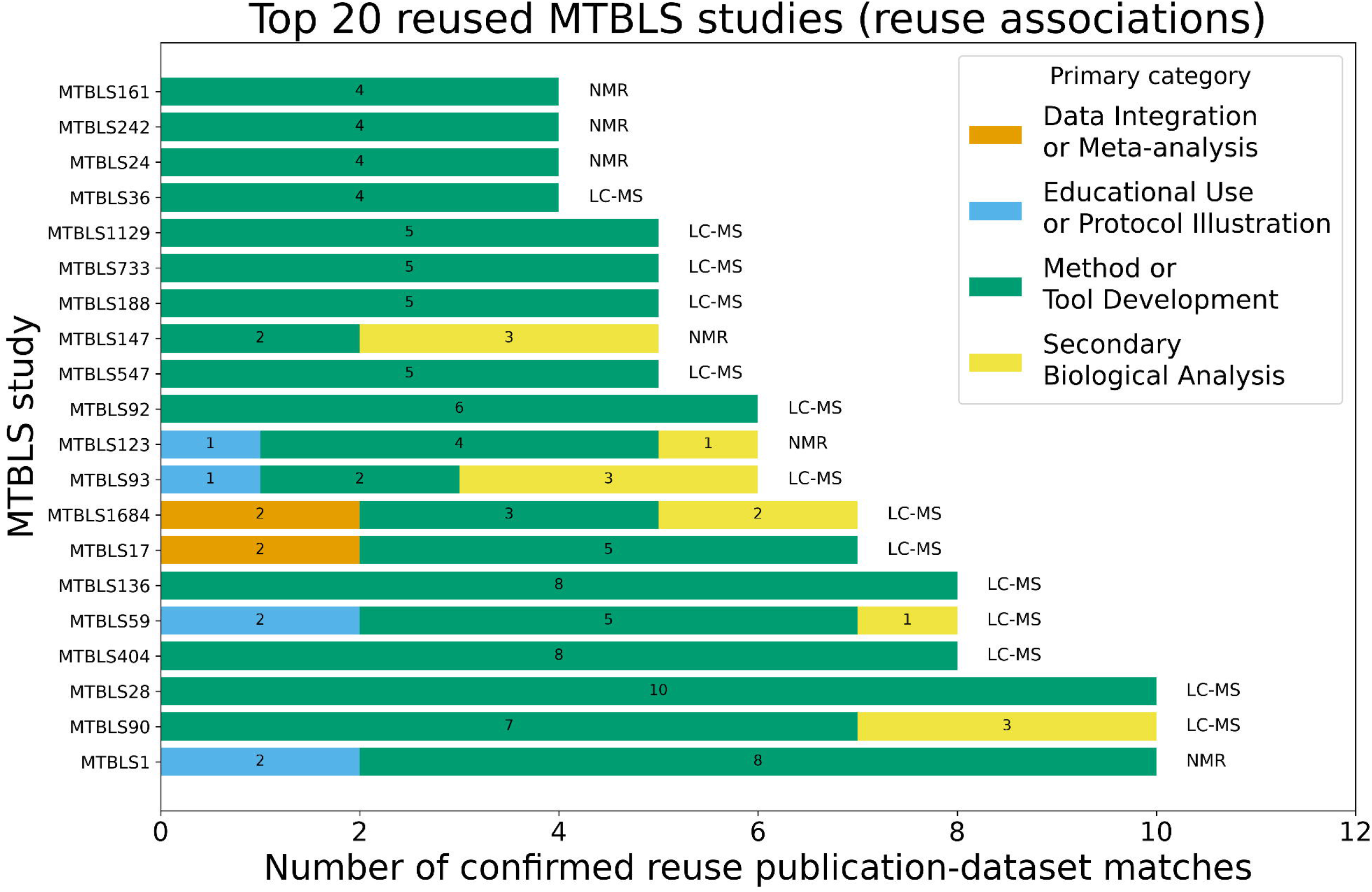
Top 20 most reused MetaboLights datasets by reuse associations, stacked by primary reuse category. Each reuse association represents one distinct confirmed publication–dataset pair.

#### 3.5.3 Data Reuse Across Release Cohorts

Figure 6 shows the release years of the 20 most reused MetaboLights datasets, which span 2012-2020. Figure 6b groups the same datasets by release cohort (year) and shows that datasets from 2017-2020 appear alongside the earliest datasets. Highly reused datasets were released across several years; this descriptive comparison does not test whether dataset age affects reuse.

**Fig 6.**
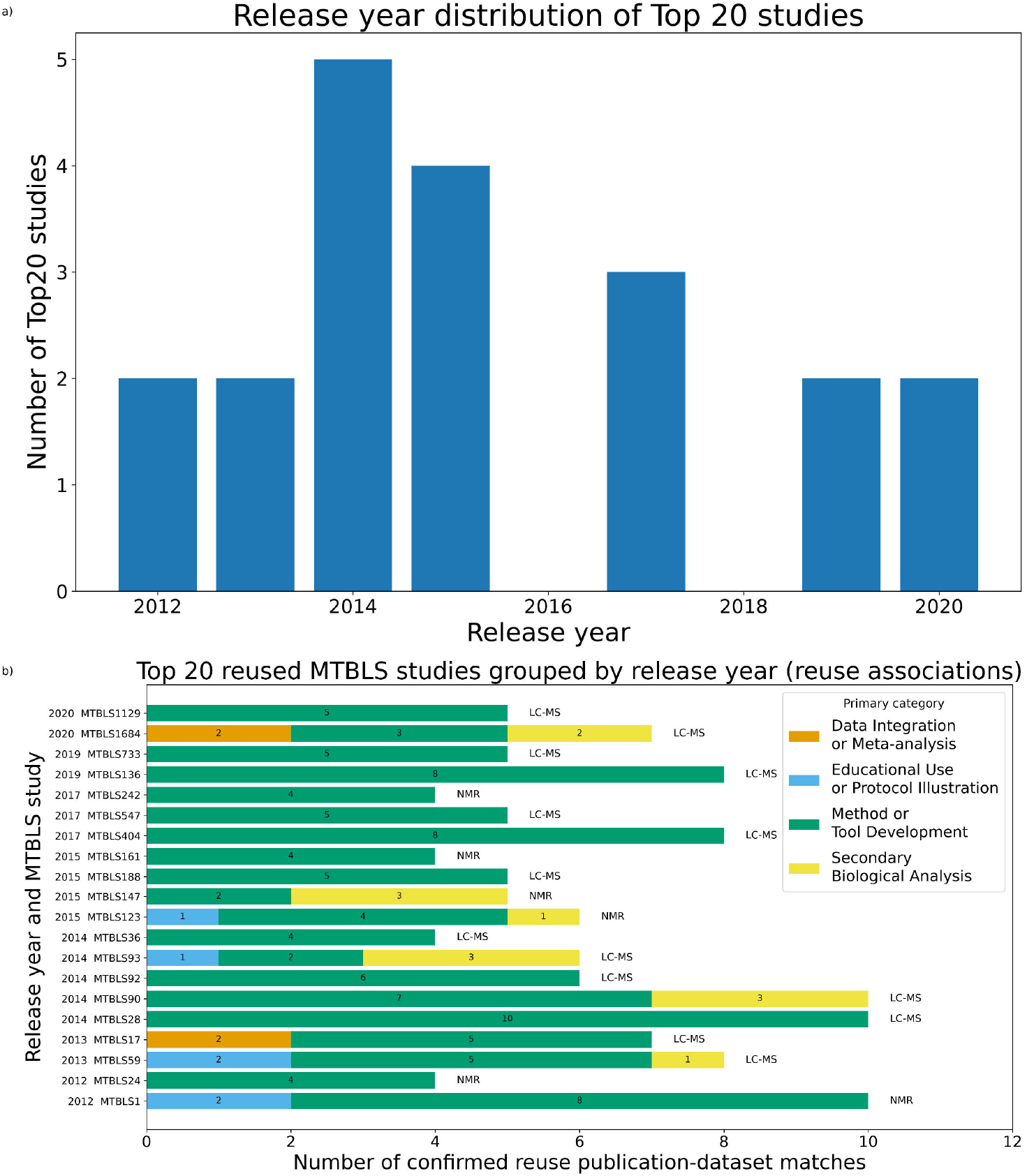
Release-year distribution of the top 20 MetaboLights datasets by reuse associations (a) and the same datasets grouped and sorted by release year (b). Each reuse association represents one distinct confirmed publication–dataset pair.

#### 3.5.4 Characteristics of the Top 5 Datasets

Table 4 describes the five MTBLS datasets with the most confirmed reuse publications. All five are human-sample studies with varied study designs. The table reports study attributes recorded in MetaboLights and reuse profiles summarised from the reviewed publications; we did not assess statistical power or metadata completeness, or test whether any characteristic is associated with reuse frequency.

**Table 4.** Key characteristics of the top 5 most reused MTBLS datasets.

| **Dataset (Release Year)** | **Confirmed reuse publications** | **Modality** | **Sample matrix** | **Study Design** | **Reuse Profile** | **Reference publication** |
|---|---|---|---|---|---|---|
| MTBLS1 (2012) | 10 | NMR | Human Urine | Case-control | Method development; NMR benchmarking | (Salek et al., 2007) |
| MTBLS90 (2014) | 10 | LC-MS (+) | Human Blood Serum | Longitudinal population | Statistical validation; large-scale modelling | (Ganna et al., 2014, 2015) |
| MTBLS28 (2014) | 10 | LC-MS (+/-) | Human Urine | Case-control | Biomarker benchmarking; classification | (Mathé et al., 2014) |
| MTBLS404 (2017) | 8 | LC-MS (-) | Human Urine | Population cohort | Workflow benchmarking; W4M training dataset | (Thévenot et al., 2015) |
| MTBLS136 (2019) | 8 | LC-MS (+/-) | Human Blood Serum | Case-control | Cohort modelling; epidemiological analysis | (Stevens et al., 2018) |

### 3.5.5 Publication Sources

Confirmed data reuse publications were distributed across 129 publication sources (canonical title harmonisation), including journals, books, and preprint servers. The highest numbers of confirmed MetaboLights reuse publications appeared in *Metabolites* and *Metabolomics* (17 each), followed by *Analytical Chemistry* and *Nature Communications* (14 each; Table 5).

**Table 5.** Journals with >5 confirmed MetaboLights dataset reuse publications.

| **Journal** | **Number of Unique Confirmed Reuse Publications** |
|---|---|
| Metabolites | 17 |
| Metabolomics | 17 |
| Analytical Chemistry | 14 |
| Nature Communications | 14 |
| BMC Bioinformatics | 12 |
| Scientific Reports | 11 |
| Journal of Proteome Research | 8 |
| Bioinformatics | 8 |

Multidisciplinary journals such as Nature Communications and Scientific Reports also appear among the retrieved publications. However, journal counts reflect coverage and searchable full text in the five literature sources, as well as explicit identifier reporting, and are therefore not an unbiased measure of community-wide use.

## 4 Discussion

This study provides a conservative, manually-validated assessment of confirmed MetaboLights dataset reuse within the limits of identifier-based detection (MTBLS#). By restricting inclusion to active data reuse instances and excluding citation-only mentions, the analysis focuses on explicitly traceable reuse events at the repository level.

The time-based analysis shows an increase in the cumulative number of confirmed reuse publications from 2014 to 2025. However, because the analysis does not normalize counts by the number or age of available MTBLS datasets, source coverage, or identifier-citation practices, it cannot estimate changes in reuse propensity or attribute the observed cumulative increase to a particular cause.

Within identifier-linked publications, Method or Tool Development is the predominant observed reuse mode and LC-MS is the most frequent modality. These distributions are descriptive and do not establish preferential reuse by category or modality.

Future work will need denominator-adjusted analyses to determine whether categories or modalities differ in reuse propensity. Differences in assay coverage, sample preparation, instrumentation, identification confidence, and quantitative comparability may complicate cross-study biological integration. This study did not measure these factors and therefore cannot determine their contribution to the observed distribution of reuse categories.

Reuse is highly asymmetric at the dataset level: most dataset accessions appear in one reuse publication, while a small subset appears repeatedly. The release-year comparison shows that highly reused datasets span several years. This study did not assess data quality, metadata completeness, or experimental characteristics as predictors of reuse and therefore does not identify the reasons for recurrent reuse.

Within the Data Integration or Meta-analysis primary category, Multi-Omics Integration is the largest sub-category, accounting for 31 of 37 confirmed reuse publications. This label describes integration across different omics layers, which can occur within a single study and does not necessarily involve synthesis across independent cohorts. Its prominence should therefore not be interpreted as evidence of frequent cross-cohort meta-analysis.

Several limitations in this study should be noted. Data reuse detection in this study relies on the presence of explicit dataset accessions (e.g. MTBLS#) within the content searchable through the selected literature sources. As a result, reuse cases in which data were accessed and used without citing the corresponding dataset accession are not captured. Reuse mediated through external platforms, such as GNPS (Wang et al., 2016), or incorporated into machine-learning models may be missed when the resulting publications do not report the original MTBLS identifiers. Because screening and classification were performed by one reviewer, independent inter-reviewer agreement was not assessed.

The analysis is also constrained by the coverage and indexing capabilities of the literature databases used. Retrieval was limited to the coverage and searchable content provided by the five selected literature sources. While preprints indexed in PubMed Central and/or Europe PMC are included where available, preprints outside indexed repositories are not captured. Supplementary materials are not systematically searched unless included in the indexed full text, which may lead to additional missed data reuse instances.

In addition, this study captures only the data reuse that results in a publication. Data downloads and subsequent analyses that do not lead to a published output are not reported here. Unpublished activities, including exploratory analyses, educational use or protocol illustration, internal research, and analyses with negative results, represent an additional source of unmeasured data reuse.

Finally, the scope of this study is restricted to MetaboLights. The data reuse from datasets in other repositories such as Metabolomics Workbench (Sud et al., 2016), GNPS/MassIVE (Wang et al., 2016) and MetaboBank (Ara et al., 2024) have not been considered. Therefore, the patterns reported in this study should not be interpreted as representative of global metabolomics data reuse as these platforms host substantial volumes of data and support distinct reuse pathways.

## 5 Future Perspectives

Future work should distinguish complementary indicators of data use (Perez-Riverol et al., 2019). Download counts can indicate access or interest but do not demonstrate analysis; verified reanalysis could be supported by deposited derived outputs, workflow records, or links from the reused dataset to a publication; and published reuse can be established through explicit identifier-linked reporting and manual validation. Shared data dictionaries, common data elements, and clear sample and instrument summaries would make reuse and its measurement more reliable (Tenenbaum and Blach, 2018). Improved dataset citation, metadata completeness, deposition of derived resources, and reusable workflows that preserve provenance would make resulting contributions more discoverable and support future curation.

An important next step is the extension of this analysis framework to additional metabolomics repositories, including Metabolomics Workbench and GNPS/MassIVE. Preliminary data collection across these resources has identified candidate publications, but systematic, repository-specific validation and harmonisation are required to enable a direct comparison. Expanding the current methodology to a multi-repository setting would allow characterisation of global cross-repository data reuse patterns, including differences in modality coverage, community usage, and integration with computational tools and workflows.

Advances in machine learning, artificial intelligence and large-scale computational frameworks (e.g. DreaMS; Bushuiev et al., 2025) further highlight the need for mechanisms that preserve links between derived models and their underlying training datasets. Developing infrastructure to capture such reuse will be essential for accurately assessing the full impact of public metabolomics resources.

Beyond technical improvements, recognising data reuse as a measurable research output may provide an additional incentive for data sharing. Improved visibility and attribution of reuse could enhance the perceived value of public datasets and encourage more consistent and high-quality data deposition.

## 6 Acknowledgements

We thank all data submitters who made their datasets publicly available through MetaboLights and MetabolomicsHub partners for insightful discussions.

## 8 Author contribution statement

IK, TP and JAV conceptualized the content of the manuscript. IK performed research, analyzed data, and prepared the draft of the manuscript. JAV acquired funding. All authors then reviewed, edited, finalized and approved the final version of the manuscript.

## 9 Compliance with Ethical Standards

This article does not contain any studies with human and/or animal participants performed by any of the authors.

## 10 Data and Code Availability Statement

The code and the filtered January 2026 search, reviewed, and derived publication-dataset records supporting this study will be publicly available through https://github.com/EBI-Metabolights/metabolights-data-reuse and archived with a persistent DOI at https://doi.org/10.5281/zenodo.23765512.

