## Supplementary Information for "Revealing the Hidden Landscape of Public Metabolomics Data Reuse in MetaboLights"

**Supplementary Data S1.** Detailed records and summary results of MetaboLights data reuse. This Excel workbook contains 20 worksheets covering confirmed dataset–publication associations, publication details and classifications, screening outcomes, and summary results by reuse category, analytical modality, scientific domain, publication year, publication source and dataset. The df_confirmed_mtbls_matches worksheet provides the complete list of 469 confirmed associations involving 273 datasets and 274 publications. The publication_table worksheet summarises confirmed publications, while screening_outcomes documents the final screening decisions for all 884 publications.

**Supplementary Table S1.** Manually-assigned reuse sub-categories.

| Secondary Biological Analysis | Method or Tool Development | Data Integration or Meta-analysis | Educational Use or Protocol Illustration |
| --- | --- | --- | --- |
| Biomarker Discovery/Signature | AI/Deep Learning | Cross-Platform Harmonization | Best Practices/Guidelines |
| Clinical Translation/Stratification | Alignment/RT/Batch | Cross-Study Meta-analysis | FAIR/Data Management |
| Differential/Case-Control Analysis | Analysis Workflow | Multi-Omics Integration | Protocol/SOP |
| Dose-Response/Exposure/Toxicology | Benchmarking/Comparative Evaluation | Network-based Integration | Teaching Dataset/Case Study |
| Environmental/Epidemiology | Classification/Regression | Systematic Review/Evidence Synthesis | Tutorial/Workshop/Vignette |
| Longitudinal/Time-Series | Cloud/Scalability/Reproducibility |  |  |
| Pathway/Network Biology | Data Structures/Formats |  |  |
| Replication/Validation | Databases/APIs (including knowledge bases and knowledge graphs) |  |  |
|  | FAIR/Data Management |  |  |
|  | Feature Selection/Dimension Reduction |  |  |
|  | Identification/Annotation |  |  |
|  | Identifier/Ontology Mapping |  |  |
|  | Meta-Analysis |  |  |
|  | Machine Learning |  |  |
|  | Network-based Integration |  |  |
|  | Normalization/Imputation |  |  |
|  | Pathway/Network Tools |  |  |
|  | Power/Sample Size Calculation |  |  |
|  | Preprocessing/Deconvolution |  |  |
|  | Visualization/Reporting/QC |  |  |
|  | Workflow/Orchestration |  |  |

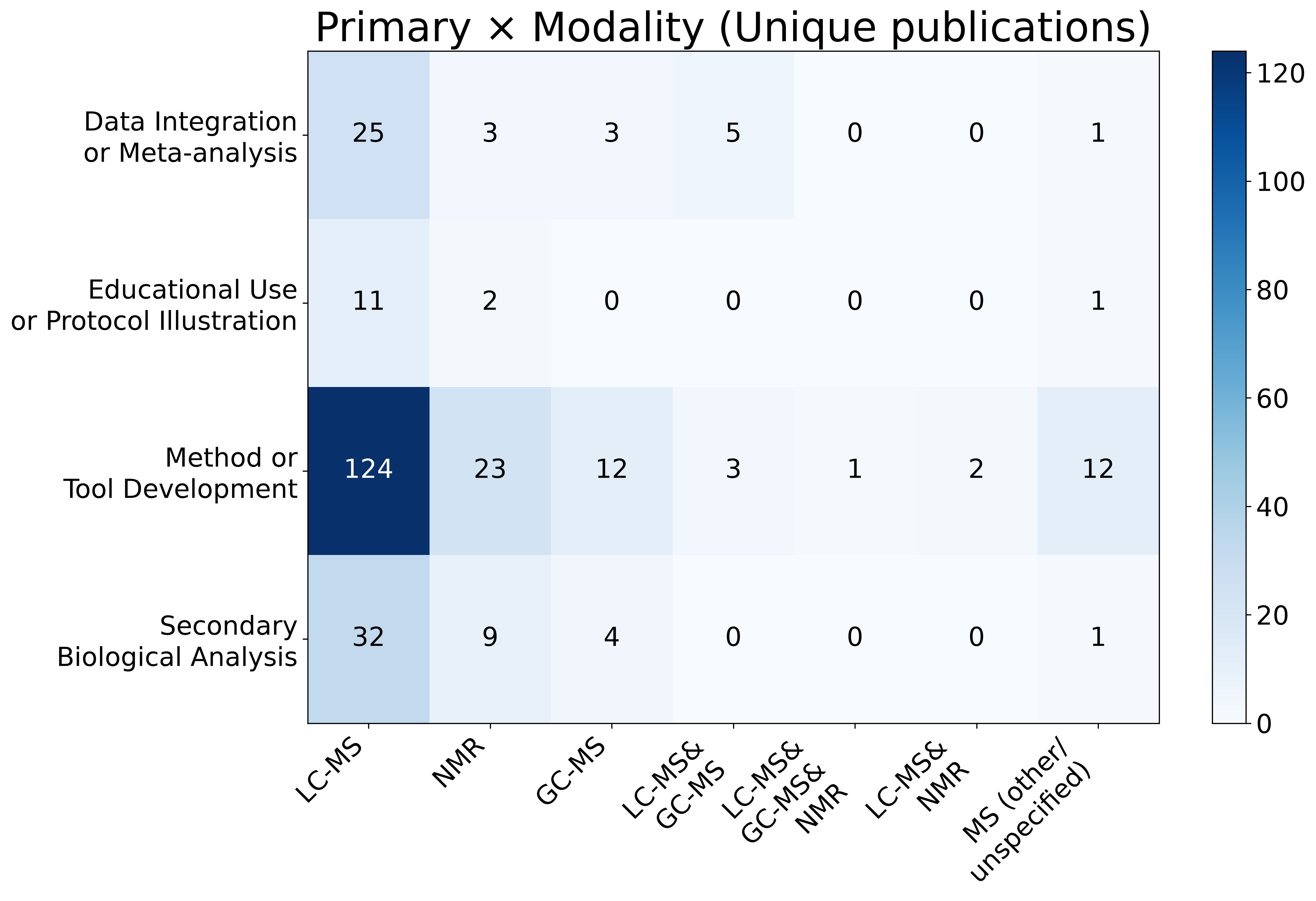

**Supplementary Figure S1.** Heatmap of unique reuse publications by primary category and analytical modality.

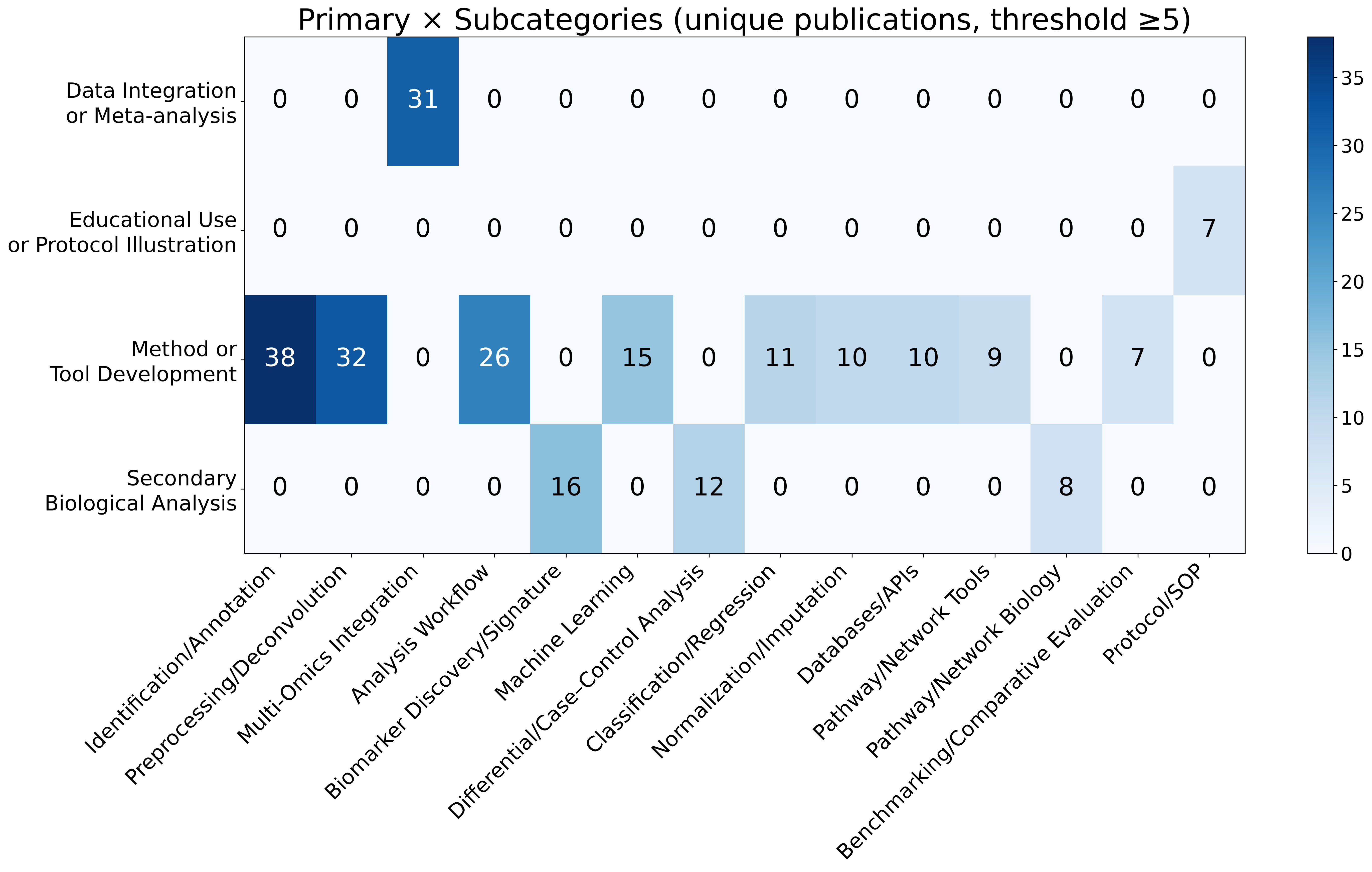

**Supplementary Figure S2.** Heatmap of unique reuse publications by primary category and sub-category (threshold ≥ 5). Sub-categories with fewer than 5 publications are omitted for clarity.

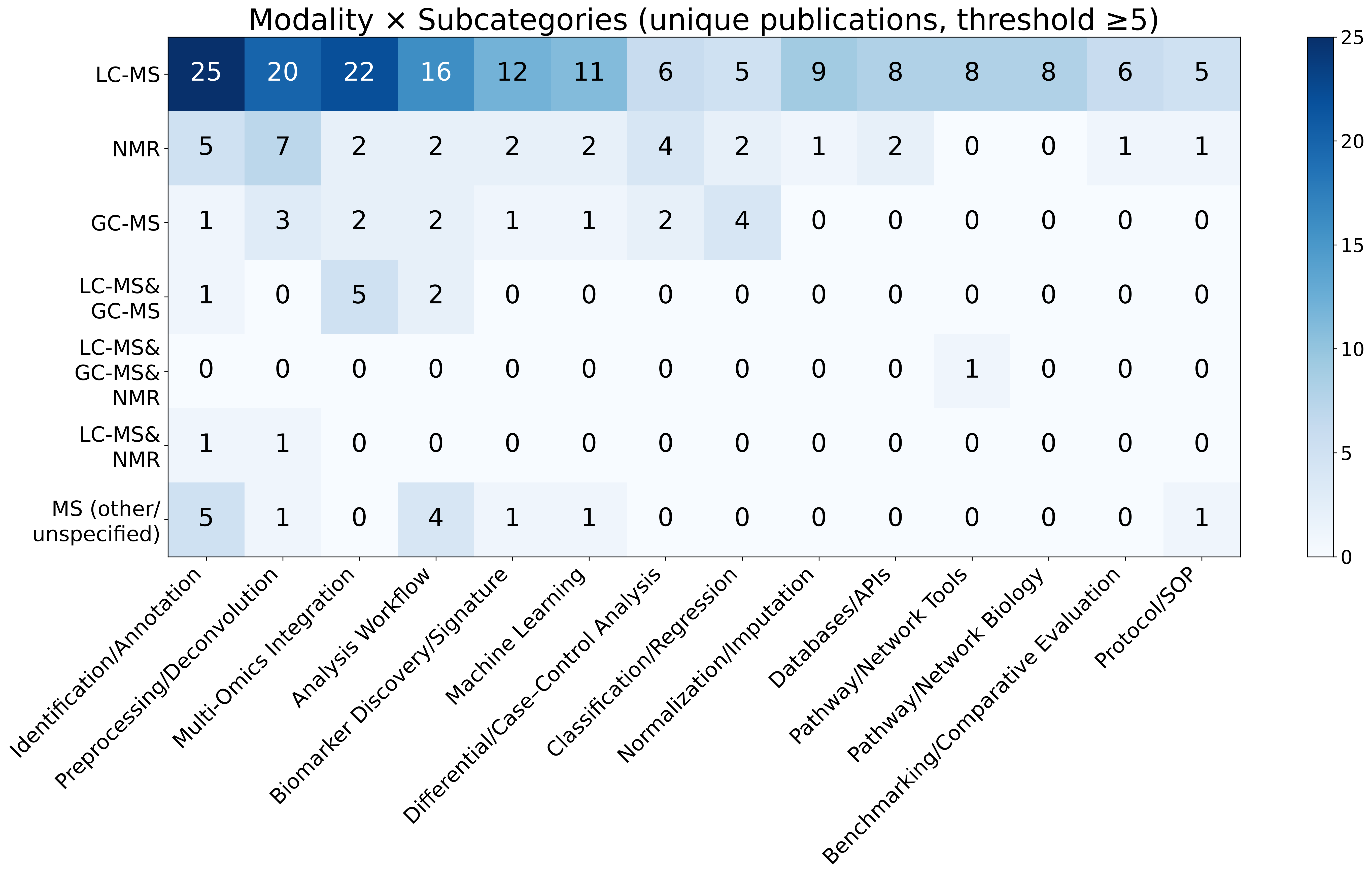

**Supplementary Figure S3.** Heatmap of unique reuse publications by analytical modality and sub-category (threshold ≥ 5).
